# Analysis of root volatiles and functional characterization of a root-specific germacrene A synthase in *Artemisia pallens*

**DOI:** 10.1101/2023.11.01.564656

**Authors:** N.R. Kiran, Ananth Krishna Narayanan, Soumyajit Mohapatra, Priyanka Gupta, Dinesh A. Nagegowda

**Affiliations:** Molecular Plant Biology and Biotechnology Lab, CSIR-CIMAP Research Centre, Bengaluru – 560065, India; Biotechnology Division, CSIR-Central Institute of Medicinal and Aromatic Plants, Lucknow-226015, India; Academy of Scientific and Innovative Research (AcSIR), Ghaziabad-201002, India

**Keywords:** *Artemisia pallens*, davana, root terpenoids, terpene synthase, germacrene A synthase

## Abstract

Davana (*Artemisia pallens*) is a valuable aromatic herb within the Asteraceae family, highly prized for its essential oil (EO) produced in the aerial parts. However, the root volatiles and their specific composition, and genes responsible for root volatiles have remained unexplored until now. Here, we show that *A. pallens* roots possess distinct oil bodies and yields ∼0.05% of EO, which is primarily composed of sesquiterpenes β-elemene, neryl isovalerate, β-selinene, and α-selinene, and trace amounts of monoterpenes β-myrcene, D-limonene. This shows that, besides aerial parts, roots of davana can also be a source of unique EO. Moreover, we functionally characterized a terpene synthase (ApTPS1) that exhibited high *in silico* expression in the root transcriptome. The recombinant ApTPS1 showed the formation of β-elemene and germacrene A with *E,E*-farnesyl diphosphate (FPP) as a substrate. Further, detailed analysis of assay products revealed that β-elemene was the thermal rearrangement product of germacrene A. Furthermore, the functional expression of ApTPS1 in *Saccharomyces cerevisiae* confirmed the *in vivo* β-elemene/germacrene A synthase activity of ApTPS1. At the transcript level, *ApTPS1* displayed predominant expression in root, with significantly lower level of expression in other tissues. This expression pattern of *ApTPS1* positively correlated with the tissue-specific accumulation level of β-elemene. Overall, these findings provide fundamental insights into the EO profile of davana roots, and the contribution of ApTPS1 in the formation of a major root volatile.

**Main conclusion:** The study demonstrated that *Artemisia pallens* roots can be a source of terpene-rich essential oil and root-specific ApTPS1 forms germacrene-A contributing to major root volatiles.

## Introduction

Plants produce an amazing diversity of compounds termed “specialized or secondary metabolites” as a means of protection against various abiotic and biotic stresses, attraction of pollinators and seed dispersing animals, and interaction with the surrounding environment (Nagegowda and Gupta 2020). The production and emission of volatile compounds by various plant species and their organs represent an adaptive trait enabling plants to interact with their surrounding environment thereby providing ecological advantage (Chen et al., 2011). Among the wide array of volatile compounds, terpenes constitute a prominent and diverse class of specialized metabolites that are produced and released by plants (Schwab et al. 2013). Volatile terpenes produced by plants have been shown to serve as signals for communication and interaction between plants and a range of other organisms, including insects, fungi, and bacteria (Huang et al., 2003; Huang et al., 2012; Schulz-Bohm et al., 2017). Also, they play pivotal roles in safeguarding plants against a variety of abiotic and biotic stresses (Unsicker et al. 2009; Loreto and Schnitzler 2010). The volatile terpenes produced in plants mainly comprise the isoprene (C_5_), monoterpenes (C_10_), sesquiterpenes (C_15_), and some diterpenes (C_20_) (Rosenkranz and Schnitzler 2016). In plants, terpenoids are derived from two common C_5_ precursor molecules, isopentenyl diphosphate (IPP) and its allylic isomer dimethylallyl diphosphate (DMAPP). While IPP is converted to isoprene by isoprene synthase, both IPP and DMAPP are utilized by prenyl diphosphate synthases to form geranyl diphosphate (GPP), farnesyl diphosphate (FPP), and geranylgeranyl diphosphate (GGPP) that serve as precursors for mono-, sesqui-, and di-terpenes, respectively (Nagegowda and Gupta 2020).

The reaction for the final conversion of prenyl diphosphates into terpenes is catalysed by terpene synthases (TPSs), which is a large class of enzymes with several sub families that produce structurally diverse cyclic and acyclic terpene skeleton via carbocationic intermediates, formed by divalent metal co-factor dependent removal of the diphosphate group (Chen et al., 2011; Christianson, 2006). Majority of TPSs can form multi-products from the same substrate, some can even utilise more than one substrate. In plants, expression of TPS is regulated in a stage-, tissue-, and species-specific manner suggesting a highly optimized production and deployment of distinct terpenes. Although most work on plant TPSs and their respective terpene products has focused on the phyllosphere, an increasing number of studies demonstrate their important roles in the rhizosphere. Root volatile terpenes have been suggested to influence the behaviour of herbivorous insects (Robert et al. 2012) and nematodes (Rasmann et al. 2005) and affect soil bacterial and fungal communities (Kleinheinz et al. 1999; Wenke et al. 2010). The majority of the reported root-specific TPSs and their enzymatic products play role in plant defence against soil borne pathogens and insects. For example, TPS encoding 1,8-cineole synthase is expressed in Arabidopsis roots and releases 1,8-cineole, which has been implicated in defense against microbes and insects (Chen et al., 2004). Similarly, β-caryophyllene was shown to be released from roots of *Centaurea stoebe* in a constitutive manner and function as a belowground plant-to-plant cue, adjusting the germination, growth, and defense of sympatric neighboring plants (Huang et al. 2019; Gfeller et al. 2019).

Besides their defensive and ecological roles, terpenes produced by root-specific TPSs, like terpenes produced by TPSs of aerial tissues, have commercial value to humans as EO for their aromatic or fragrant properties. The EOs produced in roots of several plant species have been exploited commercially due to their immense aromatic value. For instance, the EOs extracted from roots of vetiver (*Vetiveria zizanioides*) (Mallavarapu et al. 2012), ginger (*Zingiber officinale*) (Ekundayo et al. 1988), and spikenard (*Nardostachys jatamansi*) (Takemoto et al. 2009; Gupta et al. 2012) are rich in terpenoids and have high commercial value. *A. pallens* Wall. ex DC is an aromatic herb belonging to the family Asteraceae. It is naturally found in humid habitats in the plains all over India, and has been commercially cultivated for its highly prized EO derived from aerial parts of the plant. The EO of *A. pallens* plant is composed of sesquiterpenes that include mainly *cis*-davanone (53.0%), bicyclogermacrene (6.9%), and *trans*-ethyl cinnamate (4.9%), along with minor amounts of other sesquiterpenes and monoterpenes (Singh et al. 2021). The davana EO has commercial demand in local as well as international markets due to its importance in fragrance industries and as flavouring agent in preparation of cakes, pastries and some beverages (Kumara et al. 2023). Though the EO of davana aerial parts has been well explored and commercially exploited, no studies have been attempted to investigate the root volatile composition. Here, we showed that davana roots can be source of EO as they are composed of oil bodies that are rich in volatile terpenes. Further functional genomics resulted in characterization of a TPS that is specifically expressed in roots and involved in the production of a major terpene present in the root EO.

## Material and Methods

### Plant material

*A. pallens* plants were grown in the field or pots at CSIR-Central Institute of Medicinal and Aromatic Plants, Research Centre, Bengaluru. Leaf, flower bud, and root samples were collected from the fully grown plants. Samples were snap-frozen on the spot by liquid nitrogen, were ground into fine powder, and stored in -80 °C deep freezer until further use.

### Microscopy of davana root tissue

To find out cellular accumulation of organic volatile compounds, davana roots were observed under a compound microscope (Biocraft scientific systems Pvt. Ltd., India; Model-BLE-29). The cross-section (CS) and the longitudinal sections (LS) were dissected using a fine dissection blade, sample slides were prepared without any stains. Slides were observed under the microscope at 10X, 40X, and 100X with oil immersion.

### Identification of terpene synthase genes and phylogenetic analysis

Total RNA was isolated from the leaf and root tissues of three month old plants grown in the open field using the TRIzol method). The raw transcript sequences obtained from Illumina sequencing of root transcriptome were analysed, assembled and annotated by using Galaxy web analysis platform. Candidate transcripts were identified on the basis of their NCBI, and UniProt annotation. The transcripts were translated using ExPASy translate tool (http://web.expasy.org/translate). Multiple sequence alignment was generated using ClustalW and ESpript version 3.0. Sequence relatedness and rooted neighbor-joining phylogenetic tree with 1000 bootstrap value was generated using Molecular Evolutionary Genetics Analysis tool version 11 (MEGA 11) (Tamura et al. 2021). The accession number of protein sequences used in this study were listed in the Table S1.

### GC-MS analysis of *A. pallens* root essential oil

Three months old plants were uprooted, washed in running tap water and transferred to Clevenger unit for hydro-distillation for 5 hours. EO was collected and treated with sodium sulphate anhydrous (Na_2_ SO_4_) to remove water molecules. EO was diluted 1000 times in pentane and subjected to GC-MS analysis (Agilent Technologies 7980[A gas chromatograph system with the 5977[A mass selective detector). The HP5-MS column with dimension 30[m × 250 µm having film thickness 0.25 µm was used. Helium in a split ratio of 10:1 and flow rate of 1 ml/min was used as the carrier gas. The running condition for the samples was 60 °C for 5 min as initial hold, subsequently 150 °C at the flow rate of 3 °C/min, followed by a ramping of 10 °C/min until the temperature reaches 300 °C and final hold for 10 min after the temperature reaches 300 °C with a ramp rate of 10 °C/min. Mass spectrometry was conducted at 230 °C as a transfer line and ion source temperature while, 150 °C as quadrupole temperature, 70[eV ionization potential and 50 to 550 atomic mass units scan range. Version 2.0 g of NIST/EPA/NIH mass spectral library (2011/2017) was used for compound identification (Agilent Technologies, Palo Alto, CA, USA). The relative abundance of particular constituent was considered as the area percent.

### Relative quantification of β-elemene (germacrene A) in different tissues

To investigate the relative accumulation of the germacrene A, equal amount of tissues (100 mg) were taken from different parts of the plant and volatiles were extracted by grinding the tissue in liquid nitrogen followed by incubating the tissue in hexane (1 mL) overnight at 37 °C. Geraniol was added to the hexane as internal standard. The hexane extract was treated with charcoal and sodium sulphate anhydrous to remove pigments and traces of water, respectively, before using for analysis by gas chromatography fitted with a flame ionization detector (GC-FID). Mean, standard error and number of replicates were used for statistical evaluation using GraphPad QuickCalc online software (http://www.graphpad.com/quickcalcs/ttest1.cfm). The statistical significance of differences between different tissue samples was tested by unpaired Student’s *t*-test. The minimum gene-expressing tissue was used as a baseline to do the calculations.

### Protein expression and purification

Total RNA was isolated from leaf, flower and root tissues and cDNA was constructed as described before. Forward primer 5′-<u>GGATCC</u>ATGGACATGCTCGAAGCAGACA-3′ and reverse primer 5′-<u>GAGCTC</u>TTAATATTGCATGGGGATAGAAGTCTCG -3′ were used to amplify full-length ORF of the gene. This amplicon was cloned into pJET1.2 blunt vector and the correctness of the sequence was verified by Sanger’s sequencing. The sequence confirmed gene was then sub-cloned into the *Bam*HI and *Sac*I sites of pET28a bacterial expression vector, in-frame of the C-terminal (His)_6_-tag (Novagen, http://www.emdbiosciences.com) resulting in pET28a:ApTPS1. For recombinant protein expression, pET28a:ApTPS1 and pET28a (control) constructs were transformed into *E. coli* BL21 Rosetta-2 competent cells and induced when OD_600_ was 0.4 by supplementing with 0.1 mM isopropylthio-β-galactoside (IPTG) and grown in an incubator shaker at 18 °C with speed of 180 rpm for 16 hrs. The recombinant protein was partially purified by affinity chromatography on nickel-nitrilotriacetic acid (Ni-NTA) agarose beads (Qiagen, http://www.qiagen.com). Purification steps were carried out as per the manufacturer’s instructions. The purified protein was subjected to SDS-PAGE and stained with Coomassie brilliant blue. The protein concentration was determined by the Bradford method (Bradford 1976) of protein quantification.

### Terpene synthase assay and product identification

Terpene synthase assay was performed as described previously (Meena et al., 2017; Dwivedi et al., 2022) with minor changes. The reaction was performed in 500 µL of assay buffer (30 mM HEPES, pH 7.5, 10% [v/v] glycerol, 5 mM DTT, and 5 mM MgCl_2_) containing purified protein (50 to 100 µg) along with 50 µM FPP, the reaction was immediately overlaid with 500 µL of hexane and incubated for 3 hrs at 30 °C in a test tube, purified protein of *E. coli* transformed with EV served as control. After incubation, the upper hexane layer was transferred to GC vials (Sodium sulphate anhydrous used when it is necessary). The hexane extract was concentrated to around 100-150 µl before being subjected to GC-MS or GC analysis. Hexane extract from plant and *in vitro* biochemical assay products were initially analysed in GC (Agilent Technologies, 7890B and 5977A) using HP-5 MS column (0.25 mm diameter, 30 m long, and 0.25 μM film thickness). Samples were injected at 250 °C inlet temperature and an initial temperature of 60 °C with a split ratio of 2:1 and flow rate of 1[ml/min, hydrogen and zero air were used as carrier gas. The oven temperature was increased at the rate of 3 °C min from 60 °C to 150 °C (ramp-1) with a 10 min isothermal hold at 150 °C, then increased to 300 °C at the rate of 10 °C per min (ramp-2). To analyse at a lower temperature sample was injected into GC at 150 °C inlet temperature with the same parameters (Dwivedi et al. 2021). Same samples were analysed in GC-MS with the parameter as described earlier.

### Quantitative real-time PCR analysis

To analyse gene expression, reverse transcriptase quantitative polymerase chain reaction (RT-qPCR) was employed using the SYBR Green PCR Master Mix (Takara, Dalian, China) in a 48-well plate and analysed using a StepOne Real-Time PCR system (PE Applied Biosystems, http://www.appliedbiosystems.com). Total RNA isolated by TRIzol method of RNA isolation, it is treated with 10 U of DNaseI prior to use, two micrograms of RNA was used for cDNA synthesis as per manufacturer’s instructions (High-Capacity cDNA Reverse Transcription Kit was from Applied biosystems, Waltham, USA). The real-time primers (Table S2) were designed by using Primer Express software (Applied Biosystems, Waltham, USA) at the 5′ region of the gene. The experiment was performed as described by (Kumar et al. 2020) and actin served as endogenous controls in RT-qPCR. The real-time gene expression analysis of ApTPS1 and the housekeeping gene was performed and the parameters as follows: Initial denaturation 94 °C for 10 min, followed by 40 cycles of 94 °C for 15 s and 60 °C for 1 min. Fold change differences in gene expression were analysed using the comparative cycle threshold (*Ct*) method. Relative quantification was carried out by calculating *Ct* to determine the fold difference in gene expression [Δ*Ct* target – Δ*Ct* calibrator]. The relative level was determined as 2^-ΔΔCT^ (Rao et al. 2013).

### Yeast transformation and sesquiterpene production

To express the recombinant ApTPS1 protein in yeast, the ORF was PCR amplified using high-fidelity DNA polymerase using gene specific primers Table S2. Amplicon was cloned into pJET1.2 blunt vector and sub cloned into *Not* I and *Sac* I sites of pESC-leu2d expression vector, resulting in pESC-ApTPS1 construct. The resulting construct was transformed into yeast strain AM94 (doi:10.1186/1475-2859-11-162) for the production of sesquiterpenes. Yeast cultures were grown in YPD medium (1% yeast extract, 2% peptone and 2% glucose), selection of transformed yeast was done in synthetic defined (SD) medium (0.67% yeast nitrogen base with ammonium sulphate, 2% glucose and supplemented with the appropriate amino acids). Agar agar (2%) was added in case of solid medium. Yeast was grown at 30 °C, while liquid cultures were grown in an orbital shaker set to 180 rpm. Transformed yeast where genes cloned under GAL1/10 promoters were induced for expression in induction medium (SD, where glucose was substituted for galactose). Yeast cultures were overlaid with 10% dodecane (Sigma Aldrich, Cat No. D221104) and grown for 6 days at 30 °C, with 180 rpm orbital shaking. After the fermentation, the dodecane layer was collected for analysis. Analysis of sesquiterpenes produced by yeast was performed in GC using the following oven programme: initial temperature 40 °C; final temperature 220 °C; ramp-3 °C/min, total run time 60 min. Inlet was set to 220 °C and FID detector was set to 250 °C. Air flow: 250 ml/min; H_2_ fuel flow: 25 ml/min; H_2_ was used as carrier gas with flow rate of 1 ml/min. 1 µl of dodecane sample was injected in splitless mode for each analysis.

## Results

### Davana roots accumulate terpenoids in the form of oil bodies

Davana roots were carefully dissected and crushed to prepare unstained slides. These prepared slides were then examined under a compound microscope, yielding two notable observations. First, an abundance of small, spherical, and yellowish oil bodies was clearly visible (Figure 1a and 1b), appearing as distinct structures within the cell thereby providing unequivocal evidence of the presence of oil bodies within the root tissue. The second observation involved the identification of a yellowish hue within the vascular bundles of secondary and tertiary roots (Figure 1c). This fluid, had a striking resemblance in coloration with that of spherical oil bodies observed in the root cells (Figure 1). These observations indicated that *A. pallens* roots store secondary metabolites in the form oil bodies, majority of which could be volatile terpenoids. Hence, davana root tissue was subjected to hydrodistillation using a Clevenger apparatus to determine the content of oil bodies. The EO recovered after hydrodistillation was ∼0.05% of root biomass (fresh weight), and it was brick red in colour with an aromatic odour (Figure 1d). Through a comprehensive GC-MS analysis, it was found that the root oil contained 6 terpenoids which constituted major portion among the total of 37 detected compounds (Table S3). Further, among the 6 terpenoids, sesquiterpenoids constituted approximately 99% of the total composition, with neryl (*S*)-2-methylbutanoate and β-elemene/germacrene A being the predominant compounds, accounting for 23.55% and 19.61% respectively. Other sesquiterpenes like α-selinene and β-selinene were also detected accounting for 2.7% and 5.39%, respectively. There was trace amounts of monoterpenes that consisted of β-myrcene-0.16% and D-limonene-0.38% (Figure 2).

**Figure 1.**
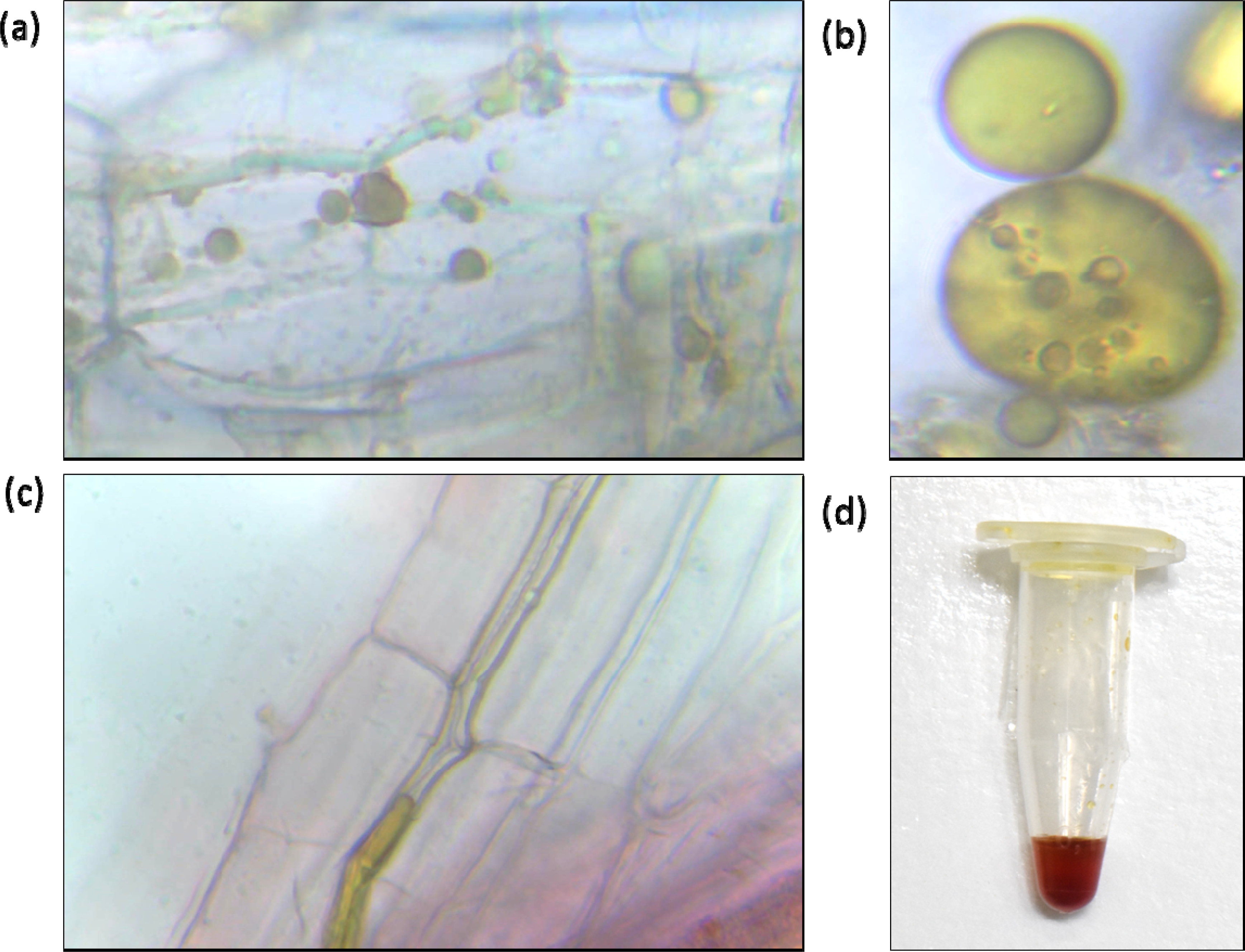
Microscopic observation of davana root tissue. (a) Davana root cell at 40X magnification showing the presence of spherical oil bodies distributed in the secondary and tertiary root cells. (b) Oil bodies in the root cells at 100X oil immersion. (c) Yellowish hue within the vascular bundles of tertiary roots (40x). (d) Essential oil extracted from the davana plant roots by hydrodistillation.

**Figure 2.**
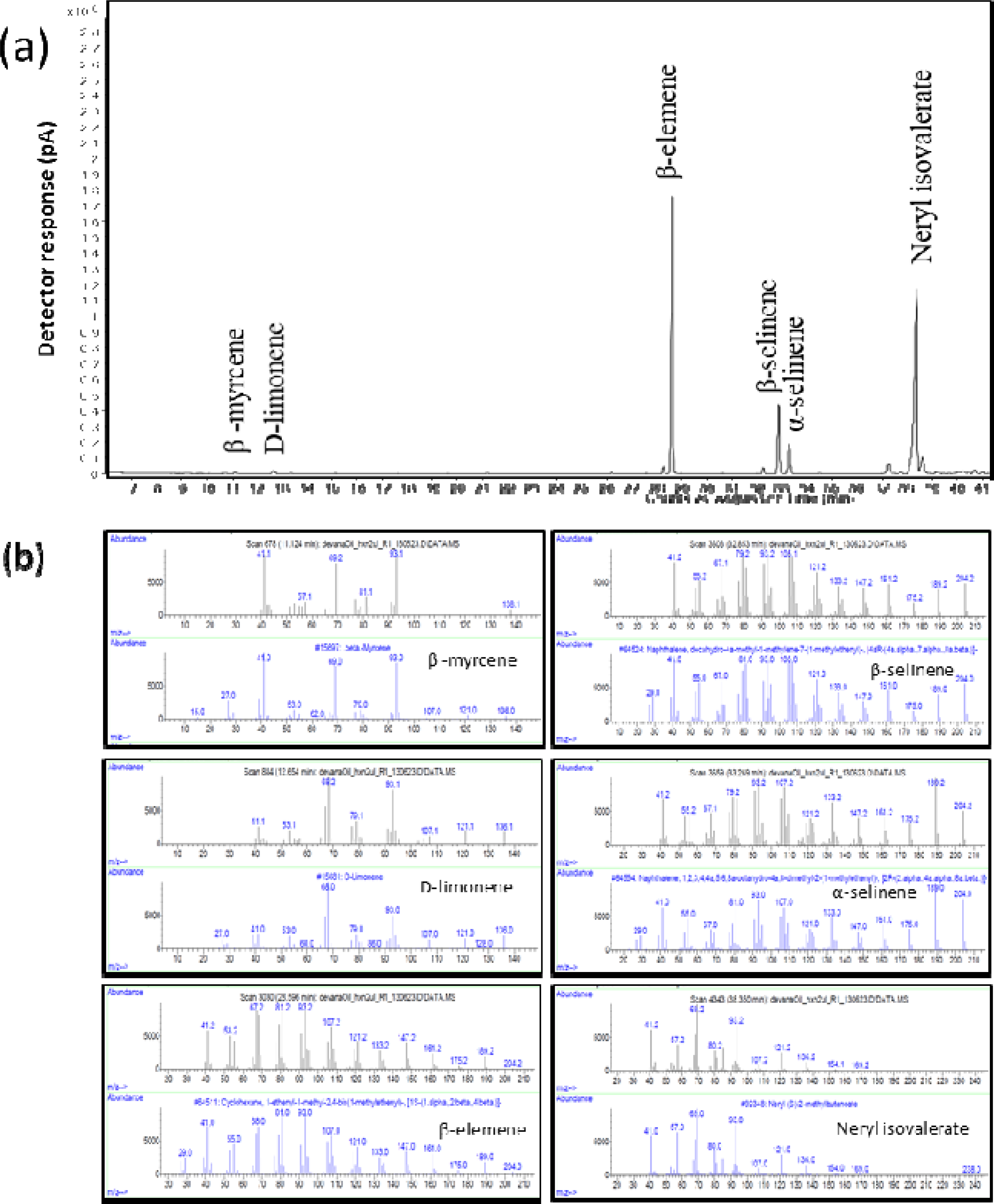
Chemical profiling of *A. pallens* root essential oil. Davana roots harvested from mature plants were subjected to hydro-distillation and the extracted essential oil was analysed by GC-MS. (a) Chromatograph showing the terpenoid components of davana root essential oil, consisting of neryl isovalerate, β-elemene, β-selinene, α-selinene, D-limonene and β –myrcene. (b) Mass spectra of identified terpenes in the oil which are aligned with the spectra of the closest compound from NIST-2017 library.

### Isolation and phylogenetic analysis of ApTPS1 from *A. pallens* roots

In order to identify terpene synthases responsible for root volatiles, the in-house transcriptome data was searched resulting in two candidate terpene synthases. Among them, one exhibited similarity to sesquiterpene synthases and named hereafter as ApTPS1, while the other (GenBank accession number OR631199) showed 93% resemblance to the *A. annua* diterpene synthase encoding geranyllinalool synthase (Chen et al., 2021). ApTPS1 was further taken for characterization. It was found that ApTPS1 has an open reading frame (ORF) of 1680 bp encoding 559 amino acids with a calculated molecular weight of 65.02 kDa. *In silico* sequence analysis for signal peptide prediction software confirmed the absence of any N- or C-terminal signal peptides in the amino acid sequence of ApTPS1 (Table S3) indicating that the protein is cytosolic, which is a characteristic feature of most sesquiterpene synthases. To functionally characterize the identified ApTPS1, it was PCR amplified using cDNA generated from root RNA of *A. pallens*, and cloned into pJET1.2 blunt cloning vector, and sequenced. The gene sequence was deposited in the NCBI database with GenBank accession number OR631198. Analysis of ApTPS1 amino acid sequence by NCBI BLASTP showed a 90.70% sequence similarity to characterized cascarilladiene synthase from *Solidago canadensis* (AAT72931) and 61.57% similarity to germacrene A synthase of *Cichorium endivia* (AZI95570) (Figure 3). The C-terminal domain contained two metal binding motifs of the aspartate-rich DDxxD and NSE/DTE (evolved from a second aspartate-rich region) to form a consensus sequence of (N,D)D(L,I,V)x(S,T)xxxE. Compared to the highly conserved DDxxD motif, the NSE/DTE motif showed relatively lower conservation across plant terpene synthases. Both of them are involved in the attachment to the trinuclear magnesium cluster (Whittington et al. 2002). The other conserved motifs like RDR deviated from earlier reports and present as RDK at the 34 amino acids upstream to DDxxD. The GTLxEL motif was conserved as GTLEEL, and an N-terminal R(R)x8W motif was also present at the 17^th^ position as RPLANFPPSIW (Li and Fan 2011). The multiple sequence alignment of the ApTPS1 sequence with 6 characterized terpene synthases of TPS-a subfamily (Figure 3) showed a common amino acid sequence at all conserved motifs except for the GTLxEL motif at 315^th^ position, where leucine (L) is substituted by isoleucine (I).

**Figure 3.**
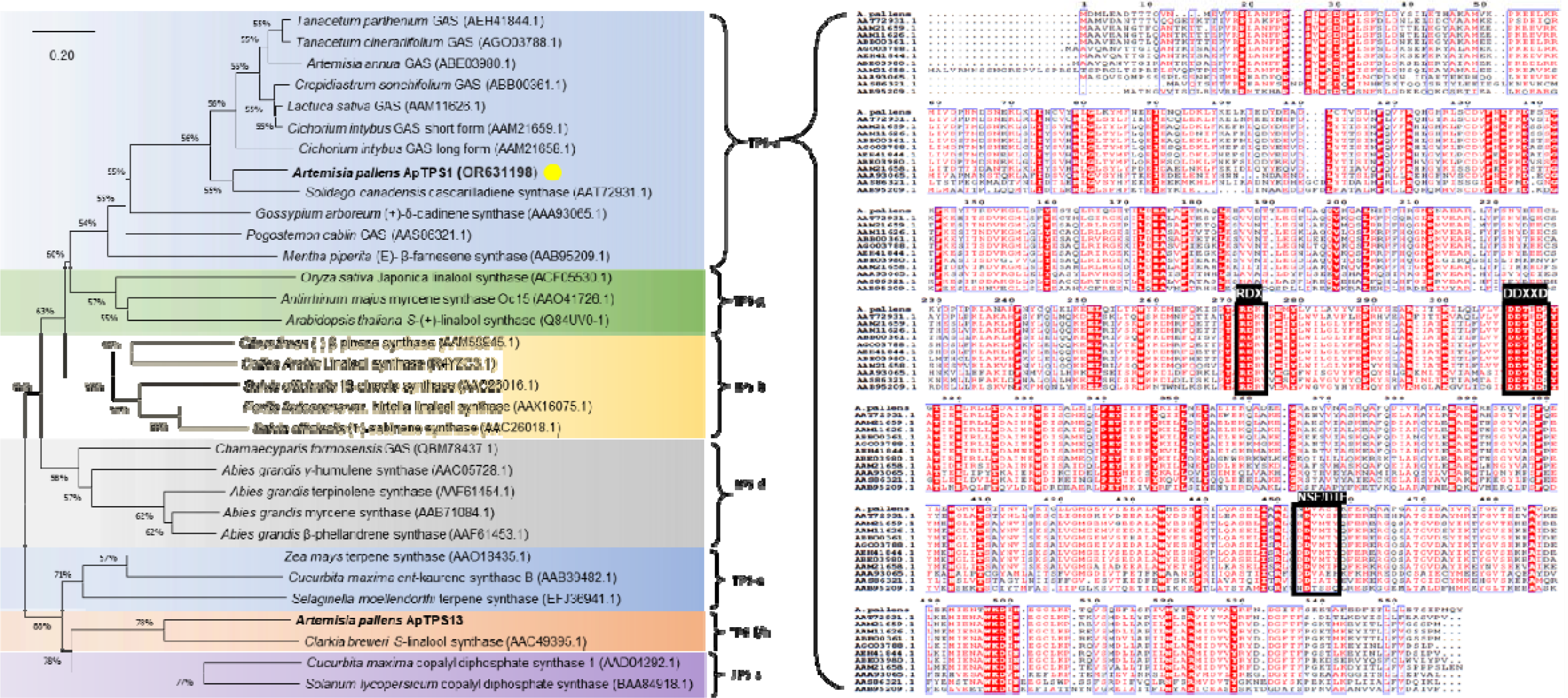
Phylogenetic analysis of *Artemisia pallens* terpene synthase 1. Phylogenetic relationship of ApTPS1 with other functionally characterized TPSs from different genera. The TPSs identified in davana root transcriptome are in bold and ApTPS1 characterized in this study is indicated by an asterisk. A phylogenetic tree was constructed by neighbour-joining method with bootstrap analysis from 1000 replicates using MEGA 11.0.11 (Tamura et al., 2007). The lengths of the branches show relative divergence among the reference ApTPS1 amino acid sequences. Tree was constructed by taking reference TPSs from different subfamilies along with identified from *Artemisia pallens* transcriptome. (a) Figure illustrating the categorization of *A. pallens* TPSs into distinct subfamilies. (b) Multiple sequence analysis of ApTPS1 with the closely related protein sequences of TPS-a subfamily. Abbreviations: GAS, Germacrene A synthase; TPS, Terpene synthase. GenBank accessions of protein sequences used in this study are listed in the Table S1.

### Functional characterization of recombinant ApTPS1

To functionally characterize ApTPS1 protein, the coding region of the gene was subcloned into a bacterial expression vector pET28a (Figure 4a) and the recombinant protein was expressed in *E. coli* and purified. Purified protein was analysed using SDS-PAGE, showing a single protein band around 65 kDa (Figure 4b). The *in vitro* assay with the purified ApTPS1 with FPP as substrate in the presence of MgCl_2_ cofactor, resulted in the formation of a single peak when analysed in GC indicating the formation of a sole product by ApTPS1. Based on the retention index analysis of the enzyme product and an authentic β-elemene standard, it was putatively predicted that the product formed by ApTPS1 is β-elemene (Figure 4c). No compounds were formed when GPP was used as a substrate, nor when MnCl_2_ and KCl_2_ were used as cofactors. Further analysis of the enzymatic product by GC-MS confirmed that the peak formed by ApTPS1 in presence of FPP corresponds to β-elemene. The mass spectra of the assay products matched with that of β-elemene standard with 91% identity and the results. It is well-known in literature that β-elemene is the thermal rearrangement product of germacrene A, via an intramolecular Cope rearrangement reaction (Reichardt et al. 1989). This has also been confirmed in our experiment by analysing the *in vitro* assay products at a lower (150 °C) and a higher (250 °C) GC inlet temperatures, where a substantial conversion of germacrene A to β-elemene was observed at higher temperatures (Figure 4c). Maximum TPS enzyme activity was observed when the assay was performed at 30 °C, however, upon increasing the incubation temperature, the product formation was drastically decreased (Figure S1a). To investigate the effect of co-factor concentration on enzyme activity, assays were performed in different concentration of MgCl_2_. A consistent rate of product formation was noted within the range of 1.5 to 10 mM MgCl_2_ concentrations. However, a decline in product formation was evident as the concentration of MgCl_2_ exceeded 10 mM (Figure S1b).

**Figure 4.**
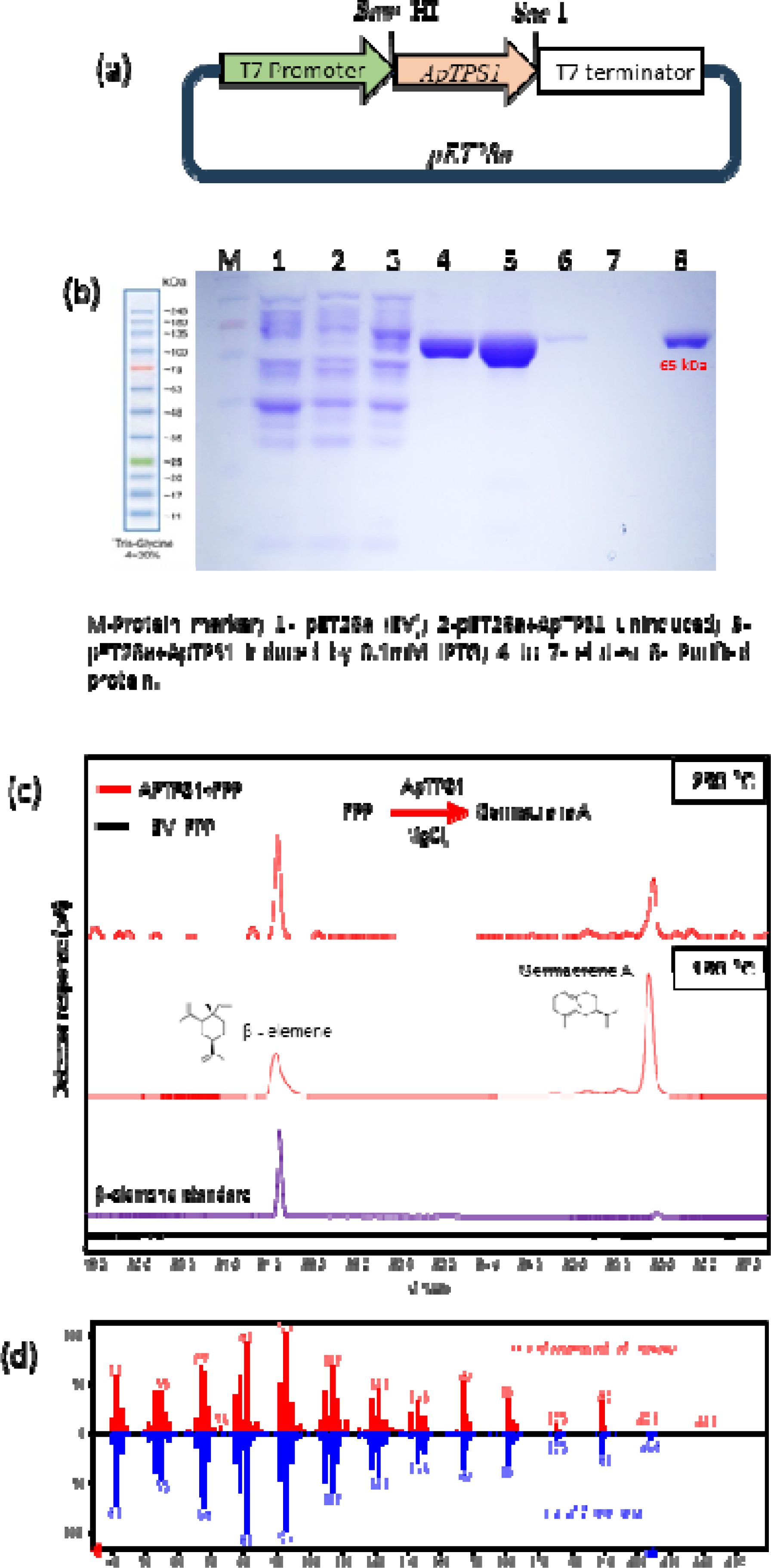
Tissue-specific *ApTPS1* gene expression and accumulation of germacrene A. (a) Analysis of transcript levels of *ApTPS1* in flower, leaf, stem and root tissues of the davana plant. Relative transcript abundance of *ApTPS1* expression in different tissues of the plant by RT-qPCR. Expression levels of *ApTPS1* were normalized to endogenous reference gene actin, and are represented as expression relative to stem that was set to 1. (b) Gas Chromatography Chromatogram showing the volatile profile of different tissues extracted by hexane, aligned with the authentic β[elemene standard. Abbreviations: ISD-internal standard.

### Expression analysis of *ApTPS1* in different tissues

To determine the expression of *ApTPS1* and the concurrent accumulation of metabolites in various tissues, quantitative real-time polymerase chain reaction (qRT-PCR) analysis using gene-specific primers, alongside relative volatile quantification was performed. The highest expression of *ApTPS1* was observed in the root (395 folds compared to stem), followed by the flower head (20 folds) and leaf (1.6-fold), while the lowest expression was noted in the stem (Figure 5a). Subsequent metabolite analysis showed the accumulation patterns of β-elemene/germacrene A correlated with the expression pattern of *ApTPS1*. This correlation between the expression of *ApTPS1* and the accumulation of β-elemene/germacrene A in plant tissues supports the active role of *ApTPS1* in *in-planta* production of β-elemene/germacrene A (Figure 5b).

**Figure 5.**
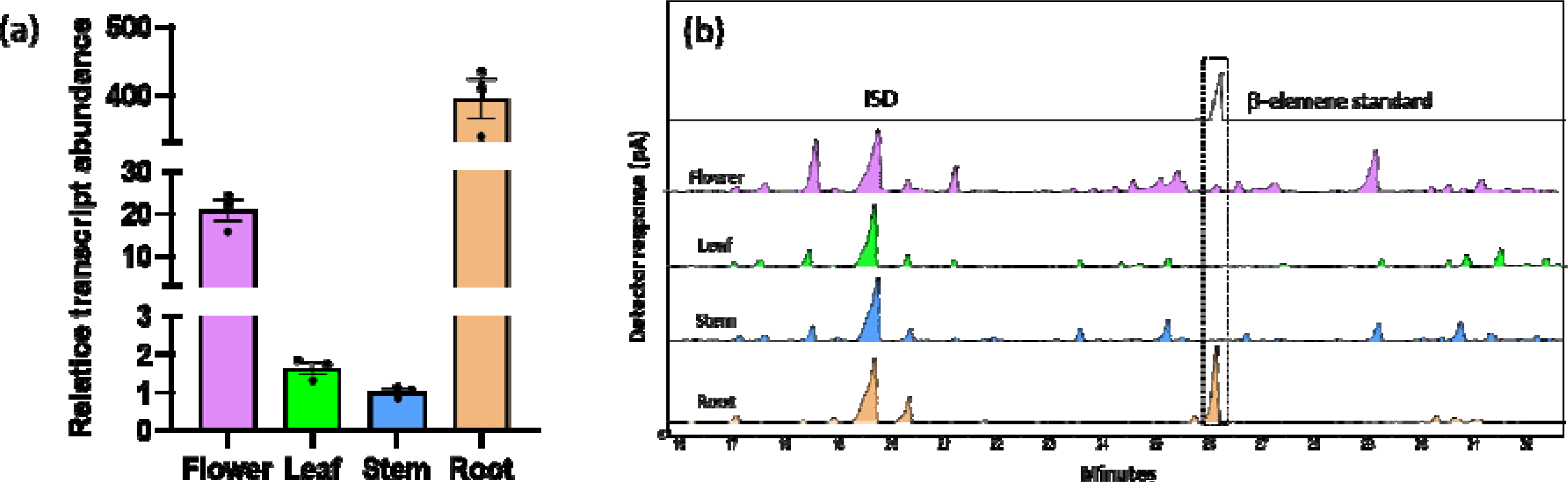
Functional characterization of recombinant ApTPS1. (a) Schematic diagram of expression cassette pET28a::*ApTPS1* for expression in *E. coli*. (b) SDS-PAGE analysis of partially purified recombinant ApTPS1. Lanes: M, protein marker; 1, total protein of empty vector transformed cells; 2, total protein of uninduced pET28a::*ApTPS1* transformed cells; 3, total protein of induced cells harbouring pET28a::*ApTPS1*; 4, elute 1; 5, elute 2; 6, elute 3; 7, elute 4; 8, desalted purified protein. (c) *In vitro* enzymatic assay product of ApTPS1 in the presence of FPP as substrate. The enzyme product was analyzed in GC at higher (250 °C) and lower (150 °C) temperatures. Purified protein from EV served as negative control. (d) Mass spectra of standard β-elemene aligned with the mass spectra of ApTPS1 enzyme assay product.

### *In vivo* functional validation of ApTPS1 in *Saccharomyces cerevisiae*

The in vitro assay using recombinant ApTPS1 revealed its role in formation of elemene/germacrene A. To further assess the *in vivo* role of ApTPS1, the ORF of *ApTPS1* was cloned into *Not*I and *Sac*I sites of pESC-leu2d yeast expression vector, resulting in generation of pESC-leu2d::ApTPS1 expression construct. The recombinant construct as well as pESC-leu2d (empty vector) were transformed into AM94 yeast strain, resulting in strain K01 and EV, respectively. GC-FID analysis of the dodecane overlay after 6 days of fermentation showed that K01 produced two unique products that were absent in the EV control (Figure 6c). Determination of the product identity using GC-MS revealed that the unique products corresponded to β-elemene (germacrene A) and Neointermedeol (Figure 6c).

**Figure 6.**
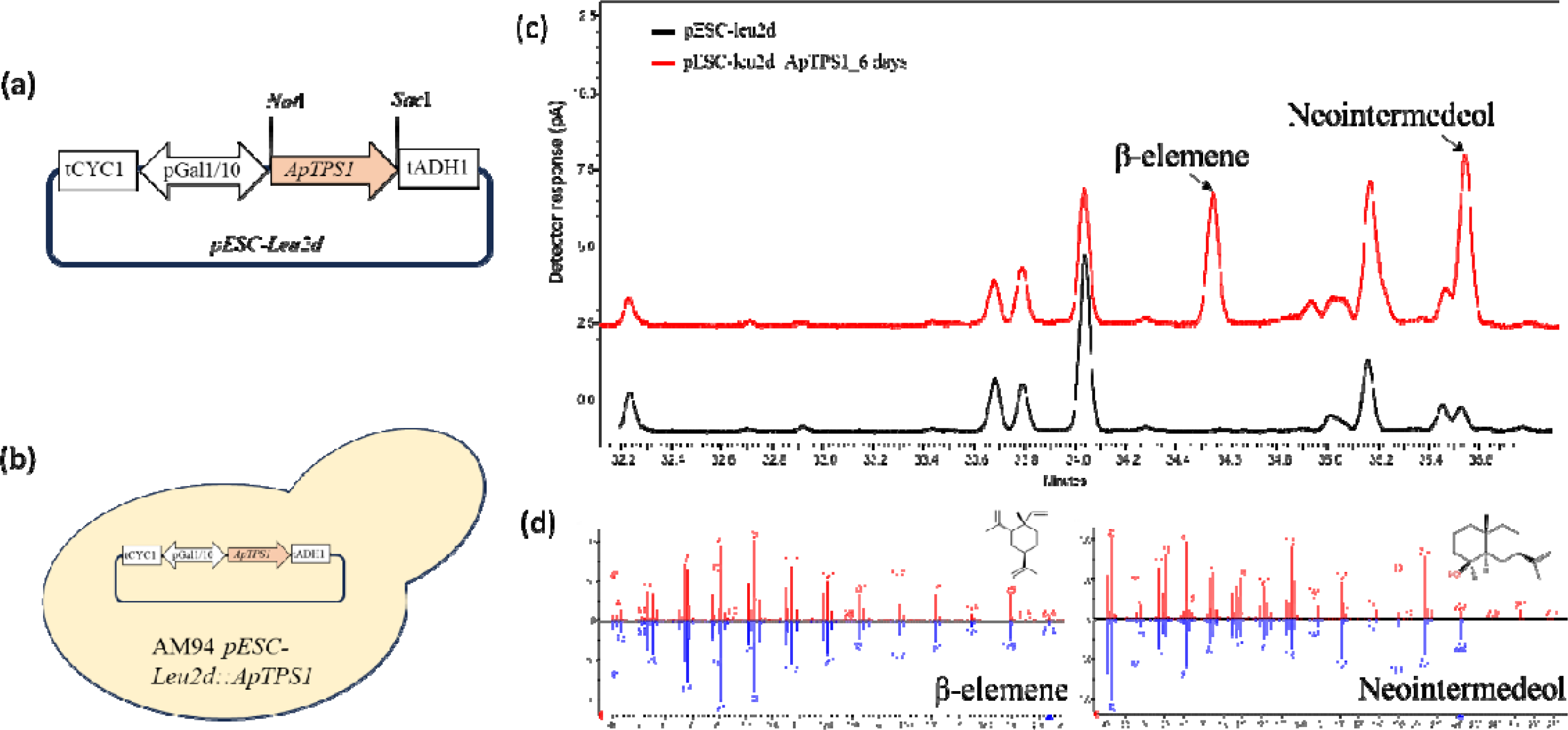
Determination of in vivo enzyme activity of ApTPS1 in yeast. (a) Schematic representation of yeast expression construct pESC-leu2d:ApTPS1. ApTPS1 was cloned into the *Not*I and *Sac*I sites of pESC-Leu2d yeast expression vector under the influence of strong Gal1-10 promoter. (b) Diagrammatic illustration showing yeast (AM94) cell transformed with pESC-leu2d:ApTPS1 construct. (c) Gas chromatograph-flame ionization detector (GC-FID) analysis of dodecane overlay containing sesquiterpenoids produced from yeast transformed with empty vector control (black line) and pESC-Leu2d:ApTPS1, after 6 days of fermentation (red line). (d) Mass spectra of β-elemene and Neointermedeol produced by recombinant yeast strain K01 aligned with the mass spectra of reference standards.

## Discussion

This work aimed to investigate the presence of EO and the corresponding genes in roots of davana (*A. pallens*), a plant known for its commercially important EO derived from its aerial parts (refs). Microscopic evaluation of Davana roots revealed the presence of oil bodies in root cells, indicating the presence of volatiles. A similar scenario involving the presence of oil bodies with a terpene composition has been reported in *Coleus forskohlii*, a species of the Lamiaceae family (Misra et al., 1994). Hydro-distillation of davana roots resulted in a sesquiterpene-rich EO (∼0.05% of fresh weight), which was composed of mainly neryl-(*S*)-2-methylbutanoate (23.55%) and β-elemene/germacrene A (∼19%) (Figure 1). The presence of acyclic sesquiterpene neryl-(*S*)-2-methylbutanoate was similar to its prominent occurrence in the roots of plants belonging to Asteraceae family such as *Achillea collina* (Kindlovits et al. 2018), *A. absinthium* and *A. vulgaris* (Blagojević et al. 2006). Whereas the second major compound of davana root EO β-elemene/germacrene A has been reported in several species in both aerial and root tissues. Among root tissue, β-elemene has been reported in *Echinops kebericho* (Hymete et al. 2007), chicory (Bouwmeester et al. 2002). Besides these major terpenoids, davana root EO also comprised of other minor sesquiterpenes such as β-selinene (5.39%), α-selinene (2.7%), lanceol (1.33%), and trace amounts of monoterpenes β-myrcene (0.16%) and D-Limonene (0.38%). Other than terpenes, the root EO also comprised of other non-terpene volatile compounds (Figure 2; Table S4).

To identify the TPSs involved in the production of davana root EO components, putative genes encoding TPSs were retrieved from the in-house generated transcriptome. This retrieval resulted in two full-length candidate TPSs, one showing homology to a sesquiterpene synthase (ApTPS1) and the other to a diterpene synthase (ApTPS13). Further investigation through *in silico* and qPCR analysis, using the candidate sesquiterpene synthase, revealed that *ApTPS1* has predominant expression in roots compared to other tissues, indicating its role in root-specific volatile formation (Figure 5). Functional characterization of the recombinant ApTPS1 and subsequent enzyme product analysis by GC-MS demonstrated that it catalyzed the formation of the cyclic sesquiterpene β-elemene from FPP. Since β-elemene is a well-known thermal cope rearrangement product of germacrene A (Reichardt et al., 1989), the enzymatic product was analyzed at a lower GC inlet temperature, which revealed a clear peak corresponding to germacrene A. This confirmed that ApTPS1 indeed formed the cyclic sesquiterpene germacrene A. It was found that ApTPS1 produced germacrene A as the sole product from FPP, with no product generated from GPP, thus indicating its bona fide germacrene A synthase (GAS) activity. Moreover, *S. cerevacea* expressing ApTPS1 also produced germacrene A, confirming the *in vivo* GAS activity of ApTPS1 (Figure 6). In the plant kingdom, numerous GASs have been identified and functionally characterized. It is noteworthy that a substantial proportion of these characterized enzymes belong to the family Asteraceae. Similar to ApTPS1, majority of GASs characterized from other plants produce germacrene A as a sole product, and no activity was found when GPP was used as substrate (Figure 4). However, there are exceptions to this scenario; for instance, GAS in yarrow (*Achillea millefolium*) formed germacrene A and selinene with FPP, and formed many monoterpenes from GPP and neryl pyrophosphate (NPP) (Pazouki et al. 2015). ApTPS1 has a calculated molecular mass of 65 kDa, which falls within the same range as that of several GAS enzymes reported in other plants. The shortest reported GAS is the *Solanum habrochaites* GAS (G8H5N2.1), consisting of 540 amino acids, while the longest GAS is the *Chamaecyparis formosensis* GAS (QBM78437.1), a gymnosperms GAS that comprises 589 amino acids (Bleeker et al., 2011; Hong et al., 2022). All three conserved motifs found in GAS were present in ApTPS1. However, in the RDR motif, the second arginine (R) amino acid is replaced by lysine (K). This kind of replacement was also observed in the closest BLASTP hit, cascarilladiene synthase, a sesquiterpene synthase from *Solidago canadensis* (AAT72931). On the other hand, the first and second aspartate-rich trinucleate metal binding motifs are completely conserved in ApTPS1 (Figure 3).

The expression pattern of the GAS gene displays adaptability in the natural world, with its expression levels varying across various plant tissues and among different plant species. For instance, in *Citrus unshiu* (Shimada et al., 2012), GAS expression was predominant in flowers, whereas in *Achillea millefolium* (Asteraceae), the highest expression of GAS was observed in both leaves and flowers (Pazouki et al., 2015). However, even within the same family, such as chicory (Asteraceae), maximal GAS expression is reported in roots (Bogdanović et al. 2020), showcasing intra-genus variability in gene expression patterns. This diversity underscores the adaptability of GAS expression to specific tissues and reflects its pivotal role in diverse biological processes within plant species. In order to determine the role of ApTPS1 in root volatile formation, a comparative analysis of gene expression and metabolite analysis in different tissues of davana was performed. This revealed a predominant production of germacrene A in roots with minor accumulation in flowers, which clearly correlated with the expression pattern of *ApTPS1* (Figure 5). This correlation of germacrene A production and ApTPS1 gene expression indicated the role of GAS (i.e., ApTPS1) in root-specific production of germacrene A in davana plant. Similar to our observation, the formation of germacrene A has been shown in roots of other plants belonging to Asteraceae family such as chicory (*Cichorium intybus*) (de Kraker et al. 1998) and *Tanacetum parthenium* L. Schulz Bip. (Majdi et al. 2011). It was shown that germacrene A undergoes a series of hydroxylation and oxidation by NADP^+^ dependent sesquiterpenoid dehydrogenase ultimately forming costunolide in chicory roots and glandular trichomes of *T. parthenium*.

Isoforms of GAS have been reported in several plant species, including Sunflower (Göpfert et al. 2010), chicory (Bogdanović et al. 2020), *Barnadesia spinosa* (Nguyen et al. 2016), and Lettuce (Bennett et al. 2002). The presence of such GAS isoforms within the same plant species highlights the complexity of terpene biosynthesis and the potential functional specialization of GAS isoforms in different biochemical pathways. In davana also, such scenario may exist as a TPS candidate having moderate *in silico* expression level in the flower, and having 70% sequence similarity to ApTPS1 was identified from the transcriptome. In general, volatiles produced from flowers or leaves are subsequently released into the external environment, attracting pollinators (Morse et al., 2012) and repelling insects (Junker & Blüthgen, 2008). In contrast, within the roots, volatile terpenoids such as germacrene A, in this case, may play an active role in plant defense mechanisms or undergo transformations into other compounds. For example, they may be converted into guianolides like lactupicrin, lactucin, and 8-deoxylactucin, which are involved in plant defense against microbes and soil-borne herbivores (de Kraker et al., 1998; Huber et al., 2016). It is reported that germacrene A acts as a central intermediate in the biosynthesis of many sesquiterpenoids (Xu & Dickschat, 2020). Given the correlation between the highest *ApTPS1* expression and the predominant presence of germacrene A in roots, it is plausible that this ApTPS1 enzyme could play a crucial role in root defense against soil-borne pathogens and insects in *A. pallens*. Overall, our study reported the terpenoid composition of the root EO of *A. pallens* for the first time. Furthermore, the study identified and characterized the GAS involved in the formation of the major volatile, germacrene A, present in davana root EO. The study also demonstrated that davana roots can serve as a valuable source of EO, in addition to floral tissues. However, the commercial value of the root EO warrants further exploration. Additionally, given the commercial significance of β-elemene/germacrene-A, the potential of ApTPS1 in synthetic biology can be further explored and harnessed.

## ACKNOWLEDGMENTS

This work was supported by the Council of Scientific and Industrial Research (CSIR) supported MLP-0003 and Aroma Mission-III. A.K.N is the recipient of Senior Research Associateship from the Indian Council of Medical Research, New Delhi. The authors express their sincere gratitude to the Director, CSIR-CIMAP for support throughout the study. The authors are also thankful to Dr. Ajit K. Shasany and Mr. Ram Krishna for helping with GC-MS analysis. The institutional communication number for this article is CIMAP/PUB/ 2023/161.

## AUTHOR CONTRIBUTIONS

N.R.K., A.K.N., S.M. and P.G. performed the experiments. N.R.K, A.K.N., S.M. and D.A.N. analysed the data. D.A.N conceived and coordinated the research. N.R.K., A.K.N. and D.A.N wrote the manuscript.

## Data Availability

All the data generated or analysed during the study are available from the corresponding author upon request.

