## Supplemental Information for "Analysis of root volatiles and functional characterization of a root-specific germacrene A synthase in *Artemisia pallens*"

**Supplementary Information:**

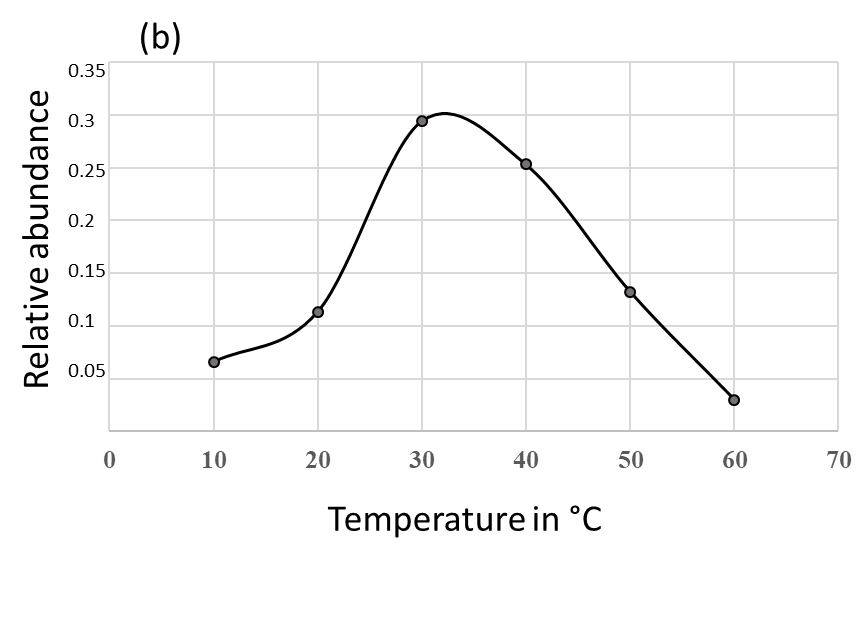

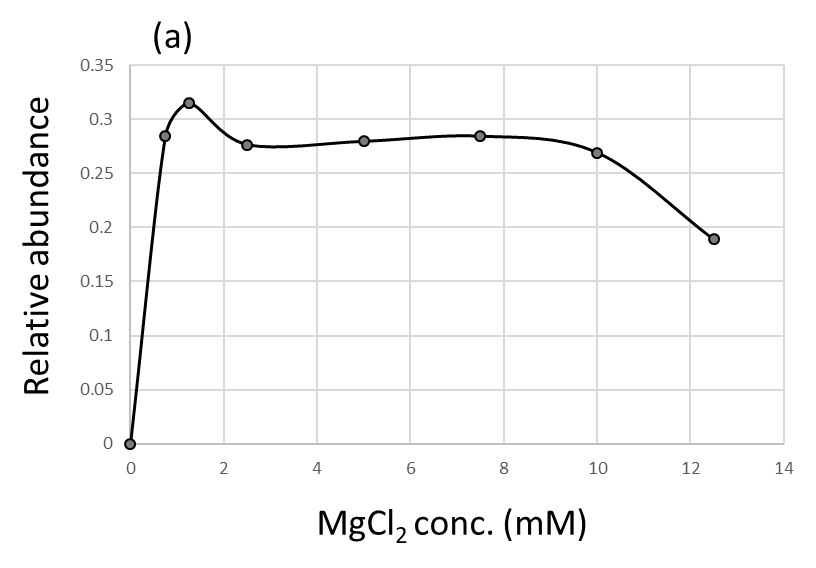

**Figure S1**: Effect of MgCl_2_ concentration on enzymatic conversion of ApTPS1 in *in vitro* assay (a). Effect of temperature on enzymatic conversion in vitro (b).

**Table S1.** Accession numbers and TPSs used in phylogenetic tree of Figure 1

| **Sl. No.** | **GenBank number** | **Description** |
| --- | --- | --- |
| 1 | OR631198 | *Artemisia pallens* GAS |
| 2 | OR631199 | *Artemisia pallens* geranyl linalool synthase |
| 2 | AAB39482.1 | *ent*-kaurene synthase B *Cucurbita maxima* |
| 3 | AAO18435.1 | Terpene synthase *Zea mays* |
| 4 | EFJ36941.1 | Terpene synthase*Selaginella moellendorffii* |
| 5 | AAC49395.1 | S-linalool synthase *Clarkia breweri* |
| 6 | BAA84918.1 | Copalyl diphosphate synthase *Solanum lycopersicum* |
| 7 | AAD04292.1 | Copalyl diphosphate synthase 1 *Cucurbita maxima* |
| 8 | AAF61453.1 | *Abies grandis* β-phellandrene synthase |
| 9 | AAB71084.1 | *Abies grandis* myrcene synthase |
| 10 | AAF61454.1 | *Abies grandis* terpinolene synthase |
| 11 | AAC05728.1 | Gamma-humulene synthase *Abies grandis* |
| 12 | QBM78437.1 | *Chamaecyparis formosensis* GAS |
| 13 | AAC26018.1 | (+)-sabinene synthase*Salvia officinalis* |
| 14 | AAX16075.1 | linalool synthase*Perilla frutescens* |
| 15 | AAC26016.1 | 1,8-cineole synthase*Salvia officinalis* |
| 16 | R4YZC3.1 | Linalool synthase*Coffea Arabic* |
| 17 | AAM53945.1 | (-)-β-pinene synthase *Citrus limon* |
| 18 | Q84UV0-1 | S-(+)-linalool synthase *Arabidopsis thaliana* |
| 19 | AAO41726.1 | Myrcene synthase Oc15 *Antirrhinum majus* |
| 20 | ACF05530.1 | Japonica linalool synthase *Oryza sativa* |
| 21 | AAB95209.1 | (E)-B-farnesene synthase *Mentha x piperita* |
| 22 | AAS86321.1 | GAS *Pogostemon cablin* |
| 23 | AAA93065.1 | (+)-δ-cadinene synthase *Gossypium arboreum* |
| 24 | AEH41844.1 | GAS *Tanacetum parthenium* |
| 25 | AAT72931.1 | Cascarilladiene synthase *Solidago canadensis* |
| 26 | AAM21658.1 | GAS long form *Cichorium intybus* |
| 27 | AAM21659.1 | GAS *Cichorium intybus* short form |
| 28 | AAM11626.1 | GAS *Lactuca sativa* |
| 29 | ABB00361.1 | GAS *Crepidiastrum sonchifolium* |
| 30 | ABE03980.1 | GAS *Artemisia annua* |
| 31 | AGO03788.1 | GAS *Tanacetum cinerariifolium* |
| 32 | AEH41844.1 | GAS *Tanacetum parthenium* |

**Table S2.** Primers used in this study along with their application

| **Sl. No.** | **Name (added restriction sites)** | **Sequence (5'-3')** | **Application** |
| --- | --- | --- | --- |
| 1 | Aptps1-FP (*Bam*H1) | GGATCCATGGACATGCTCGAAGCAGACA | ORF cloning in bacteria |
| 2 | Aptps1_RP (*Sac*1) | GAGCTCTTAATATTGCATGGGGATAGAAGTCTCG | ORF cloning in bacteria and yeast |
| 3 | Aptps1_YFP (*Not*1) | GCGGCCGCATGGACATGCTCGAAGCAGACA | ORF cloning in yeast |
| 4 | ApTPS1_RT_FP | TACAAGAGATGTGAAGGGCT | Real-time PCR |
| 5 | ApTPS1_RT_RP | TCTACAGCCTCCTTTAGTTGAGC | Real-time PCR |
| 6 | ApACT-RT-FP | GACCAACTGGGATGACATGGAG | Real-time PCR |
| 7 | ApACT-RT-RP | TCTTCTCGCGGTTAGCCTTG | Real-time PCR |
| 8 | ApTPS1_GW_F | GGGGACAAGTTTGTACAAAAAAGCAGGCTTAATGGACATGCTCGAAGCAG | Subcellular localization |
| 9 | ApTPS1_GW_R | GGGGACCACTTTGTACAAGAAAGCTGGGTTATATTGCATGGGGATAGAAGTCTCG | Subcellular localization |
| 10 | ApACT-RT-FP | GACCAACTGGGATGACATGGAG | Housekeeping gene |
| 11 | ApACT-RT-RP | TCTTCTCGCGGTTAGCCTTG | Housekeeping gene |

**Table S3.** GC-MS analysis of davana root volatile composition.

| **­­­­** | **RT (min)** | **Hit Name** | **Chemical formula** | **Quality** | **Mol Weight (amu)** | **% area** |
| --- | --- | --- | --- | --- | --- | --- |
| 1 | 11.124 | β -Myrcene | C_10_H_16_ | 64 | 136.125 | 0.16% |
| 2 | 12.654 | D-Limonene | C_10_H_16_ | 94 | 136.125 | 0.38% |
| 3 | 26.159 | 5,9,9-Trimethyl-spiro[3.5]non-5-en-1-one | C_12_H_18_O | 78 | 178.136 | 0.15% |
| 4 | 28.239 | (- ) β -elemene | C_15_H_24_ | 52 | 204.188 | 0.53% |
| 5 | 28.596 | (- ) β -elemene | C_15_H_24_ | 91 | 204.188 | 19.61% |
| 6 | 32.221 | 2-Isopropenyl-4a,8-dimethyl-1,2,3,4,4a,5,6,7-octahydronaphthalene | C_15_H_24_ | 91 | 204.188 | 0.44% |
| 7 | 32.853 | β -selinene | C_15_H_24_ | 99 | 204.188 | 5.39% |
| 8 | 33.269 | α -Selinene | C_15_H_24_ | 92 | 204.188 | 2.70% |
| 9 | 37.243 | 1-Naphthol, 5,7-dimethyl- | C_12_H_12_O | 80 | 172.089 | 1.05% |
| 10 | 37.726 | (R)-lavandulyl acetate | C_12_H_20_O | 86 | 196.146 | 0.21% |
| 11 | 38.35 | Neryl (S)-2-methylbutanoate | C_15_H_26_O_2_ | 90 | 238.193 | 23.55% |
| 12 | 38.588 | Davanone B | C_15_H_24_O_2_ | 96 | 236.178 | 2.34% |
| 13 | 39.137 | α-Selinene | C_15_H_24_ | 92 | 204.188 | 0.30% |
| 14 | 40.014 | Neryl propionate | C_13_H_22_O_2_ | 53 | 210.162 | 0.24% |
| 15 | 40.229 | Bicyclo[4.1.0]heptane, 7-(1-methylethylidene)- | C_10_H_16_ | 83 | 136.125 | 0.45% |
| 16 | 40.452 | Myrtanyl acetate | C_12_H_20_O_2_ | 64 | 196.146 | 0.25% |
| 17 | 40.675 | (+)-gamma-Gurjunene | C_15_H_24_ | 42 | 204.188 | 0.41% |
| 18 | 40.987 | α -Fenchene | C_10_H_16_ | 84 | 136.125 | 0.22% |
| 19 | 41.997 | 1,7,7-Trimethyl-2-vinylbicyclo[2.2.1]hept-2-ene | C_12_H_18_ | 25 | 162.141 | 0.15% |
| 20 | 42.391 | 4-Methyl-1,4-heptadiene | C_8_H_14_ | 46 | 110.11 | 0.23% |
| 21 | 42.77 | α -Patchoulene | C_15_H_24_ | 89 | 204.188 | 3.63% |
| 22 | 43.03 | Bicyclo[3.3.0]octan-2-one, 7-isopropylidene- | C_11_H_16_O | 49 | 164.12 | 0.48% |
| 23 | 43.387 | Lanceol, cis | C_15_H_24_O | 38 | 220.183 | 1.33% |
| 24 | 43.81 | (E)-β-Famesene | C_15_H_24_ | 43 | 204.188 | 0.13% |
| 25 | 44.59 | 1,4-Methanocycloocta[d]pyridazine, 1,4,4a,5,6,9,10,10a-octahydro-11,11-dimethyl-, (1.alpha.,4.alpha.,4a.alpha.,10a.alpha.)- |  | 27 | 204.163 | 1.00% |
| 26 | 44.791 | 3-Bromo-7-methyl-1-adamantanecarboxylic acid | C_12_H_17_BrO_2_ | 30 | 272.041 | 0.24% |
| 27 | 44.887 | Aromadendrane | C_15_H_26_ | 38 | 206.203 | 0.78% |
| 28 | 44.991 | 1,3,3-Trimethyl-2-hydroxymethyl-3,3-dimethyl-4-(3-methylbut-2-enyl)-cyclohexene | C_15_H_26_O | 43 | 222.198 | 1.83% |
| 29 | 45.132 | Pyridine-4-carboxylic acid, 1,2-dihydro-3-cyano-5,6-dimethyl-2-oxo-, methyl ester | C_10_H_10_N_2_O_3_ | 50 | 206.069 | 0.31% |
| 30 | 45.6 | (Phenylthio)acetic acid, 2,7-dimethyloct-7-en-5-yn-4-yl ester | C_18_H_22_O_2_S | 22 | 302.134 | 0.26% |
| 31 | 45.689 | Pyridine-4-carboxylic acid, 1,2-dihydro-3-cyano-5,6-dimethyl-2-oxo-, methyl ester | C_10_H_10_N_2_O_3_ | 58 | 206.069 | 2.26% |
| 32 | 45.868 | 2,5,5,8a-Tetramethyl-4-methylene-4a,5,6,7,8,8a-hexahydro-4H-chromene | C_14_H_22_O | 38 | 206.167 | 0.12% |
| 33 | 46.886 | Anthracene, 1,2,3,4,5,6,7,8-octahydro- | C_14_H_18_ | 68 | 186.141 | 1.71% |
| 34 | 48.602 | Benzenemethanol, 2-methoxy- α-2-propenyl- | C_11_H_14_O_2_ | 28 | 178.099 | 0.05% |
| 35 | 48.943 | 1,4-Naphthalenedione, 2-hydroxy-3-(1-propenyl)- | [C_13_H_10_O_3_](https://pubchem.ncbi.nlm.nih.gov/#query=C13H10O3) | 76 | 214.063 | 4.04% |
| 36 | 49.508 | 1,4-Naphthalenedione, 2-hydroxy-3-(1-propenyl)- | [C_13_H_10_O_3_](https://pubchem.ncbi.nlm.nih.gov/#query=C13H10O3) | 53 | 214.063 | 23.02% |
| 37 | 51.291 | 4,7-Dihydroxy-1,10-phenanthroline | [C_12_H_8_N_2_O_2_](https://pubchem.ncbi.nlm.nih.gov/#query=C12H8N2O2) | 64 | 212.059 | 0.07% |

**Table S4.** Prediction of subcellular targeting signals in ApTPS1 protein.

| **Prediction Program** | **Localization predicted for ApTPS1** |
| --- | --- |
| SignalP-5.0 | Cytosolic |
| TargetP - 2.0 | Cytosolic |
| WoLF PSORT | Cytosolic |
| PrediSi | Cytosolic |
| DeepLoc 2.0 | Cytosolic |
